# QRS Detection by Combinatorial Optimization With MLP Assisted Peak Scoring

**DOI:** 10.64898/2026.04.19.719501

**Authors:** Bruce Hopenfeld

## Abstract

A multiple channel QRS detector is described. The detector partitions raw signal segments into peak domains, extracts parameters associated with the peak domains, and scores peaks based on these parameters. A multi-layer perceptron (MLP) with 11 inputs generates provisional peak scores, which are refined through application of rules involving 20-30 parameters. An optimal sequence of supra threshold peaks is determined. Separately, combinatorial optimization determines an optimal structured heart rhythm sequence. Adjudication between the general supra threshold sequence and the structured sequence depends on noise level, peak quality, and rhythm structure quality. For multiple channel fusion, peak scores are determined as a noise weighted function of channel peak scores. The MLP was trained on approximately 70% of channel 1 of the MIT-BIH Arrhythmia Database. The supplementary rules were heuristically chosen over all channel 1 records. Sensitivity (SE) and positive predictive value (PPV) of the detector applied to channel 2 were a function of the noise threshold used to discard segments. At a noise level that would exclude 2.2% of channel 1 data, the SE and PPV were 99.67% and 99.75% respectively. Importantly, even in high noise, the detector was able to track large scale features of heart rhythm. Fused channel 1 and channel 2 SE and PPV were 99.96% and 99.98% respectively. The present algorithm points the way toward maximal extraction of heart rhythm information from noisy signals, and the potential to reduce false alarms generated by automated rhythm analysis software.

## 1. Introduction

QRS detection based on combinatorial optimization has been applied to single and multiple channel datasets [1-5], including the benchmark MIT-BIH Arrhythmia database (MIT-BIH DB [6,7]). This study aims to extend this previous work as follows:

- Implement combinatorial optimization based on peak scores determined in part by a neural network
- Show that combinatorial optimization enables heart rhythm tracking in relatively high noise, including cases of apparent signal dropout
- Show that a model based on a relatively small number of parameters (∼30) can produce state-of-the-art QRS detection results for the benchmark MIT-BIH DB
- Show the explicit relationship between noise measures and QRS detection capability for channel 2 of the MIT-BIH DB
- Implement multi-channel combinatorial optimization with peak score fusion that is based on noise measures
- Show that tracking the temporal relationships of peak pairs can enhance both QRS detection and, relatedly, detection of conditions such as heart block
- Provide a framework for extracting temporal and size/shape information for QRS, P, and T waves, potentially enabling further analysis of such information

## 2. Methods

### A. Single Channel Computational Framework

Figure 1 is an overview of single channel QRS detection, which is performed on short (e.g., 8 second) signal segments. A segment is partitioned into peak domains with the help of a composite signal (Csig) (e.g., blue tracing in Figure 2) that is based on first and second derivatives of the raw signal. Various peak domain parameters are extracted (see Table I), a subset (n=11) of which are input to a multi-layer perceptron (MLP). The MLP output is a provisional peak score ranging from 0-1. The scores are revised by application of approximately twenty first order, heuristically chosen rules based on various Table I parameters.

**Figure 1.**
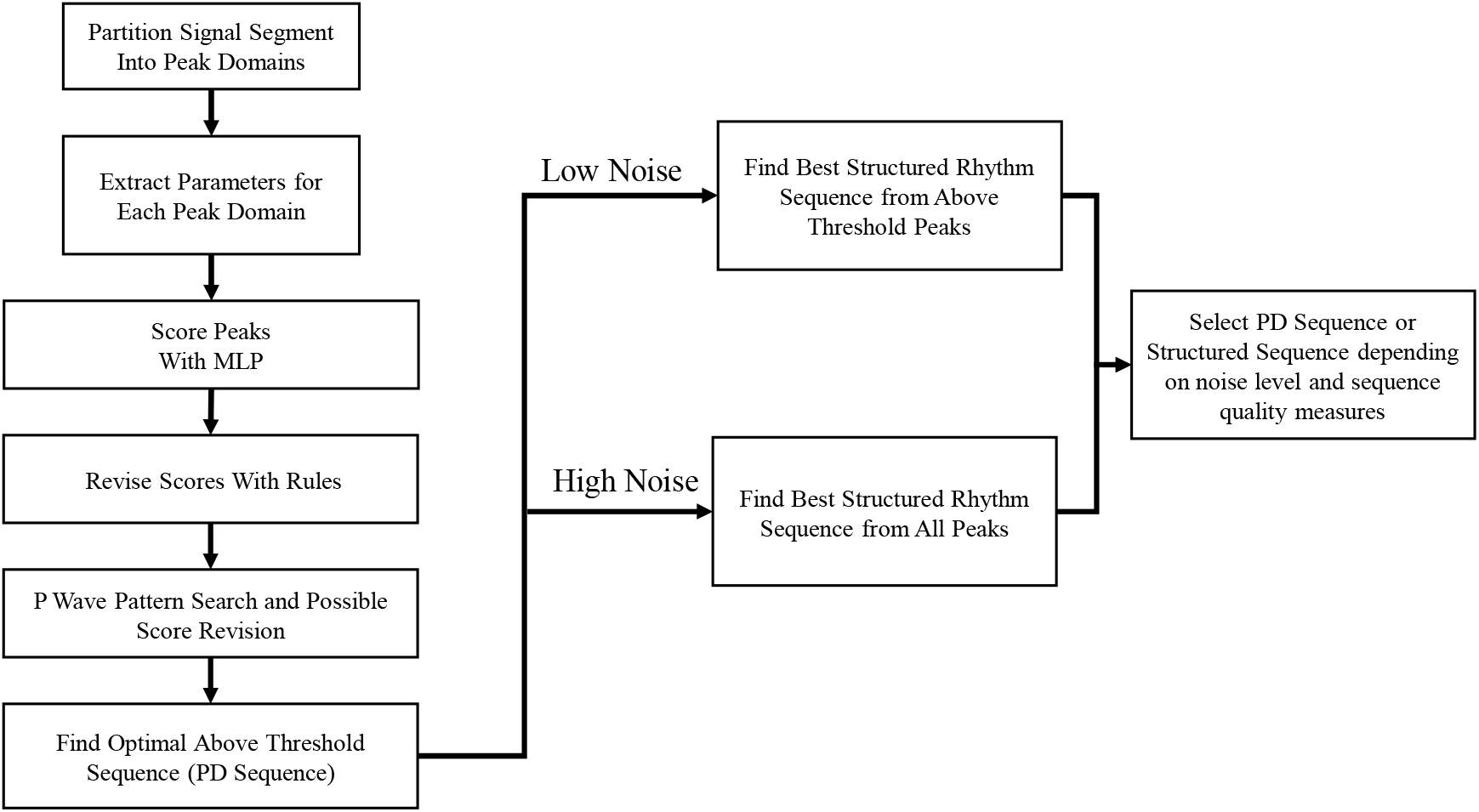

**Figure 2.**
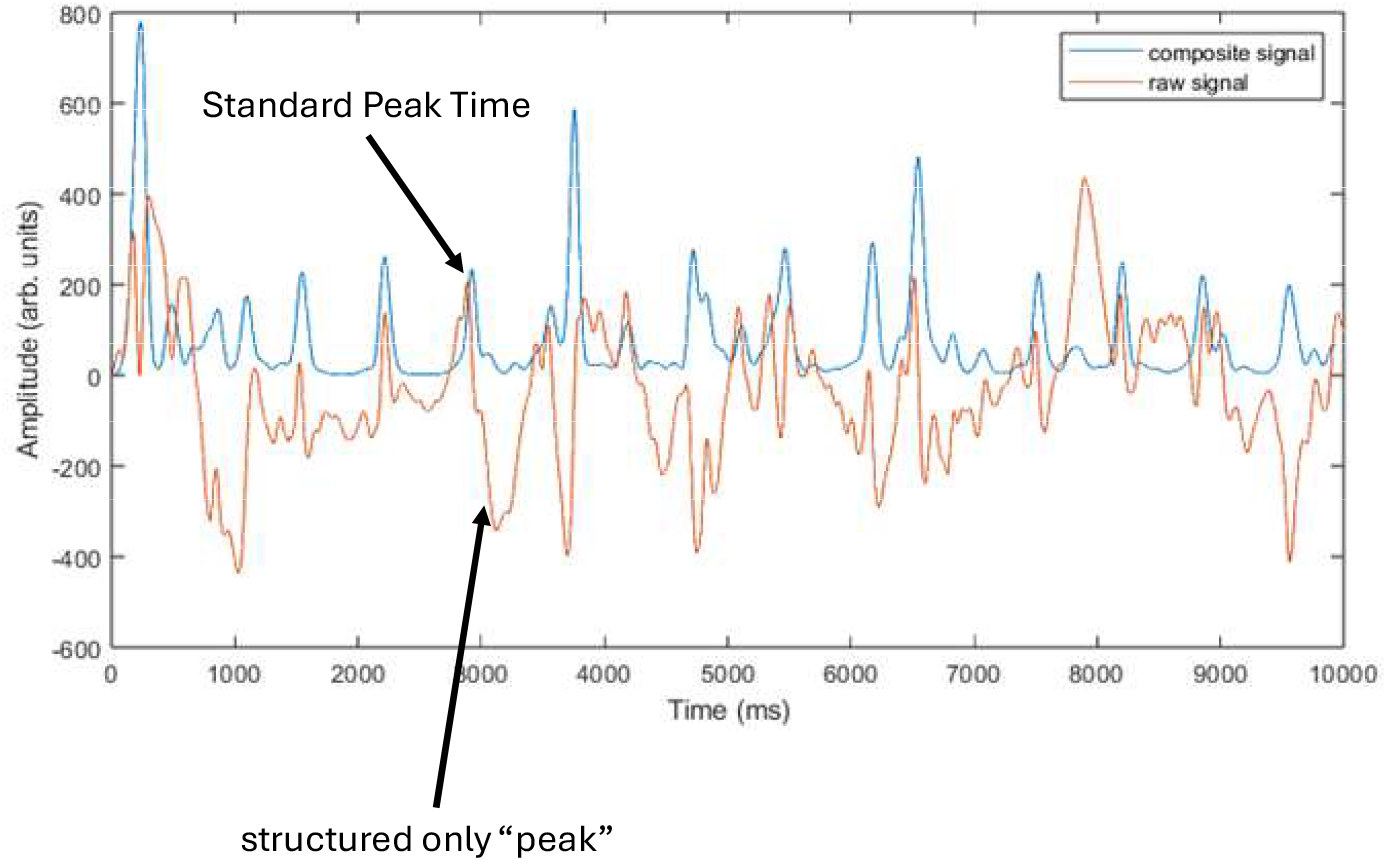

**TABLE I.**
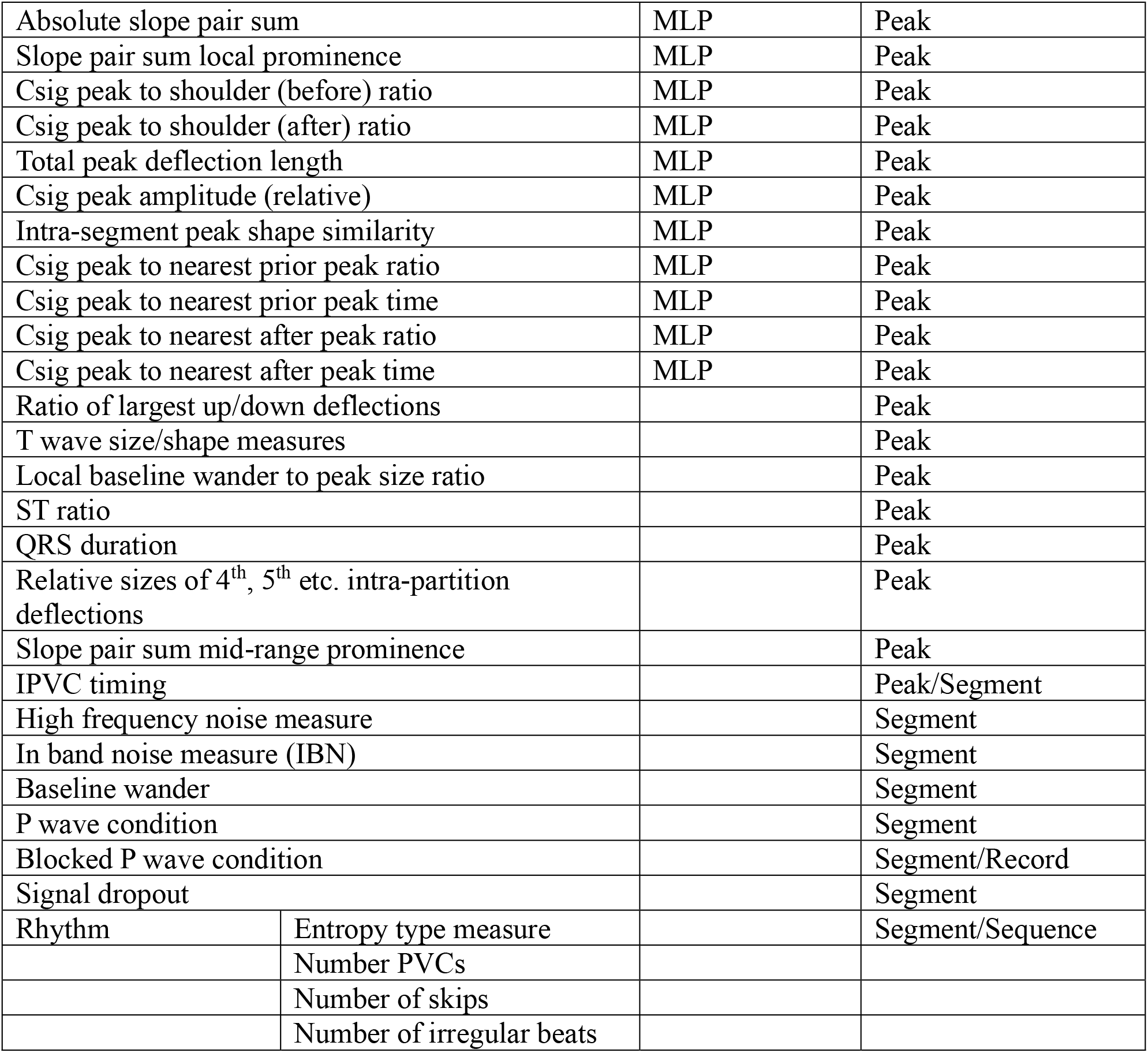

|  |  |  |  |
| --- | --- | --- | --- |
| Absolute slope pair sum |  | MLP | Peak |
| Slope pair sum local prominence |  | MLP | Peak |
| Csig peak to shoulder (before) ratio |  | MLP | Peak |
| Csig peak to shoulder (after) ratio |  | MLP | Peak |
| Total peak deflection length |  | MLP | Peak |
| Csig peak amplitude (relative) |  | MLP | Peak |
| Intra-segment peak shape similarity |  | MLP | Peak |
| Csig peak to nearest prior peak ratio |  | MLP | Peak |
| Csig peak to nearest prior peak time |  | MLP | Peak |
| Csig peak to nearest after peak ratio |  | MLP | Peak |
| Csig peak to nearest after peak time |  | MLP | Peak |
| Ratio of largest up/down deflections |  |  | Peak |
| T wave size/shape measures |  |  | Peak |
| Local baseline wander to peak size ratio |  |  | Peak |
| ST ratio |  |  | Peak |
| QRS duration |  |  | Peak |
| Relative sizes of 4<sup>th</sup>, 5<sup>th</sup> etc. intra-partition deflections |  |  | Peak |
| Slope pair sum mid-range prominence |  |  | Peak |
| IPVC timing |  |  | Peak/Segment |
| High frequency noise measure |  |  | Segment |
| In band noise measure (IBN) |  |  | Segment |
| Baseline wander |  |  | Segment |
| P wave condition |  |  | Segment |
| Blocked P wave condition |  |  | Segment/Record |
| Signal dropout |  |  | Segment |
| Rhythm | Entropy type measure |  | Segment/Sequence |
|  | Number PVCs |  |  |
|  | Number of skips |  |  |
|  | Number of irregular beats |  |  |

Next, a segment based P wave search is performed to assess whether there is a pattern of peaks with timing and amplitude characteristics of P waves and QRS complexes. If such a pattern is detected, P wave peak scores are reduced.

Peaks with scores above the detection threshold (0.5) are grouped. If all these peaks are separated from one another by at least a minimum physiological period (e.g. 220 ms), they are all tagged as provisionally detected peaks (PD Peaks). In case at least one pair is closer together than the physiological minimum, all physiologically permissible sequences are generated from the above threshold peaks and the sequence with the highest total score is selected as the PD Peaks.

A search is performed for sequences characterized by a structured heart rhythm. If noise is low, this search is performed by generating sequences from the above threshold peaks. Otherwise, a more computationally expensive search is performed across all peaks. Structured sequences are scored according to structure quality and individual peak scores. The highest scoring sequence is chosen. The PD Peak sequence is also scored for rhythm structure.

A final choice is made between the PD Sequence and the best structured sequence, based upon noise level, peak scores and the relative rhythm structure scores. In cases of very low noise, the PD Peak sequence is chosen. In cases of very high noise, the structured sequence is chosen if its structure score is sufficiently large; otherwise, the segment is considered too noisy to analyze. A noise level in between these extremes requires an adjudication of the above mentioned factors.

The MLP has a single hidden layer with 23 (=2*11+1) nodes, followed by layer normalization, ReLU, and a 2 node fully connected layer with the final output determined by a Softmax activation function.

### B. Multi-Channel Fusion

For multi-lead recordings, separate PD Peak and structured sequences are once again determined and compared. To obtain the fused scores for all peaks, temporally aligned peaks across channels are merged and assigned scores set to a noise weighted average of the individual channel scores. A peak that does not temporally align with any other channel’s peak has its score further weighted by its slope prominence over approximately one second (“Slope pair sum mid-range prominence “in Table I). This parameter serves as a local measure of signal quality. Also, if the unmatched peak corresponds to a channel that contains many P waves that are not followed by QRS complexes, the unmatched peak is rejected.

### C. Noise Assessment

Signal quality is quantified using three distinct noise metrics:

- **High-Frequency Noise:** Assessed via localized autocorrelation to identify sinusoidal type noise.
- **In-Band Noise:** Assessed by generating a low frequency baseline signal (discrete cosine transform with a 0.5Hz cutoff) and measuring the differences between this signal and QRS onsets as shown in Figure 3.
- **Baseline Wander Noise:** Based on the (curved) length of the baseline signal normalized by segment peak amplitudes.

**Figure 3.**
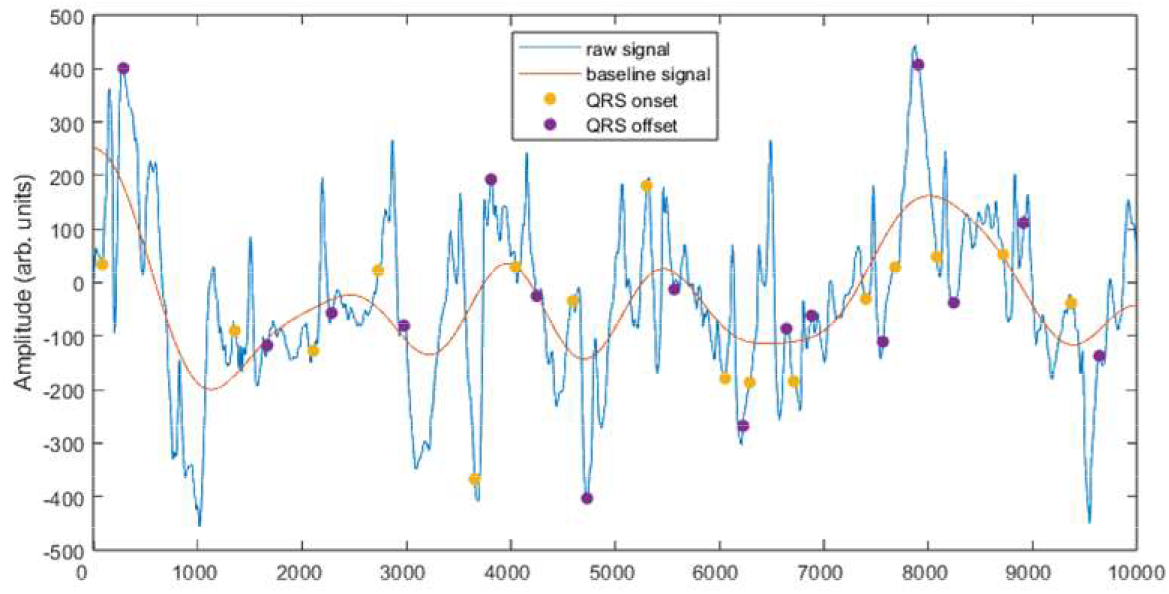

In the case of high frequency noise is detected, the signal is smoothed by a simple moving average filter. This smoothed signal is partitioned, and its peaks are scored (with a different MLP) in the same manner as the raw signal. In the case that the segment contains a large amount of high frequency noise, the raw signal is not even processed. Otherwise, in most instances, the higher of the raw and filtered scores is assigned to a peak.

The QRS onset and offsets are determined by a combination of assessing the curvature of the Csig and the first derivative of the raw signal. The onsets and offsets are not necessarily co-extensive with, and usually occur within, the boundaries of the peak partitions.

### D. Structured Sequence Regularity Scoring

The primary method for distinguishing a true rhythm from random noise is an entropy-like assessment that calculates a logarithmic penalty based on the clustering quality of the Poincare plot. A sequence with tight clusters receives a higher (less negative) score, while a random distribution of RR intervals results in a poor score. The regularity scoring handles PVCs (including pure bigeminal rhythms). However, for general structured sequence scoring, which can involve a large number of sequences, ventricular couplets (short-short-long) are ignored, which results in an artificially low regularity score for the corresponding sequence. (This limitation is due to computational costs.) The regularity score for the PD Sequence does take couplets into account.

### E. P Wave Pattern Search

A segment based P wave search is performed to assess whether there is a pattern of peaks with timing and amplitude characteristics of P waves and QRS complexes. The intervals and amplitude ratios corresponding to closely spaced peaks (consistent with the human PQ interval range) are subject to a simple clustering analysis. If the cluster quality is high, a P Wave condition is detected, and the score of the P wave type peaks is reduced accordingly. (This analysis is only undertaken only for relatively low noise segments.)

If a P Wave condition is detected, a further search is undertaken for peaks that roughly match the size and shape of the P waves, but that are not followed by larger peaks. An example is shown in Figure 4. If many such isolated peaks are detected across an entire record (e.g. Record 231, channel 2), a P Wave Heart Block condition is detected, and unmatched peaks are ignored for the purpose of multi-channel fusion, as described in Section 2.

**Figure 4.**
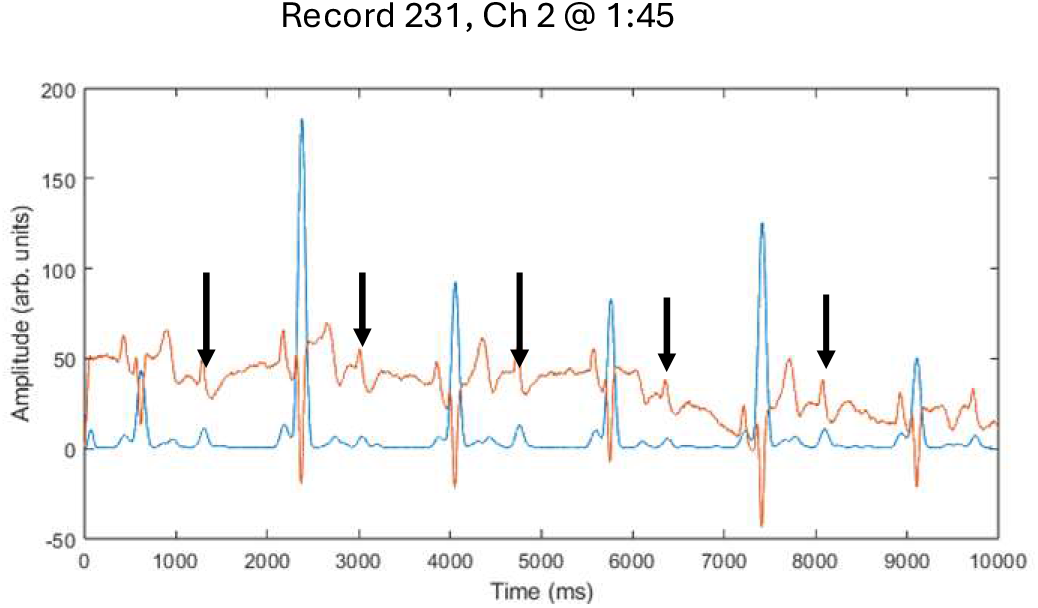

### F. Structured Sequence Only Peaks

As shown in Figure 1, noise can result in distorted QRS complexes and correspondingly long peak area partitions, which may fail to localize peaks with sufficient precision to determine sinus rhythm. To address this problem, in the case of at least moderately noisy segments, additional “structured only” peaks are designated where relatively large deflections occur temporally separated from the standard peak time, which is the apex of the composite signal. “Structured only” peaks are assigned a nominal score and may only form part of structured sequences.

### G. Dropout Detection

In some instances, signal amplitude may vary greatly over a few seconds due to e.g. electrode contact issues. An example is shown in Figure 5. Such dropouts are characterized by multi-second subsegments with very small or non-existent peaks in comparison with surrounding signal peaks. In the MIT-BIH DB, the examples of dropout exhibit a gradual baseline shift curvature, which helps to distinguish dropouts from bradycardia. The no-peak/baseline curved signal segment is isolated, differenced (to remove the baseline drift), and then subject to partition, scoring and structured sequence search. Structured sequences must be integrated with peaks from the non-dropout signal areas.

**Figure 5.**
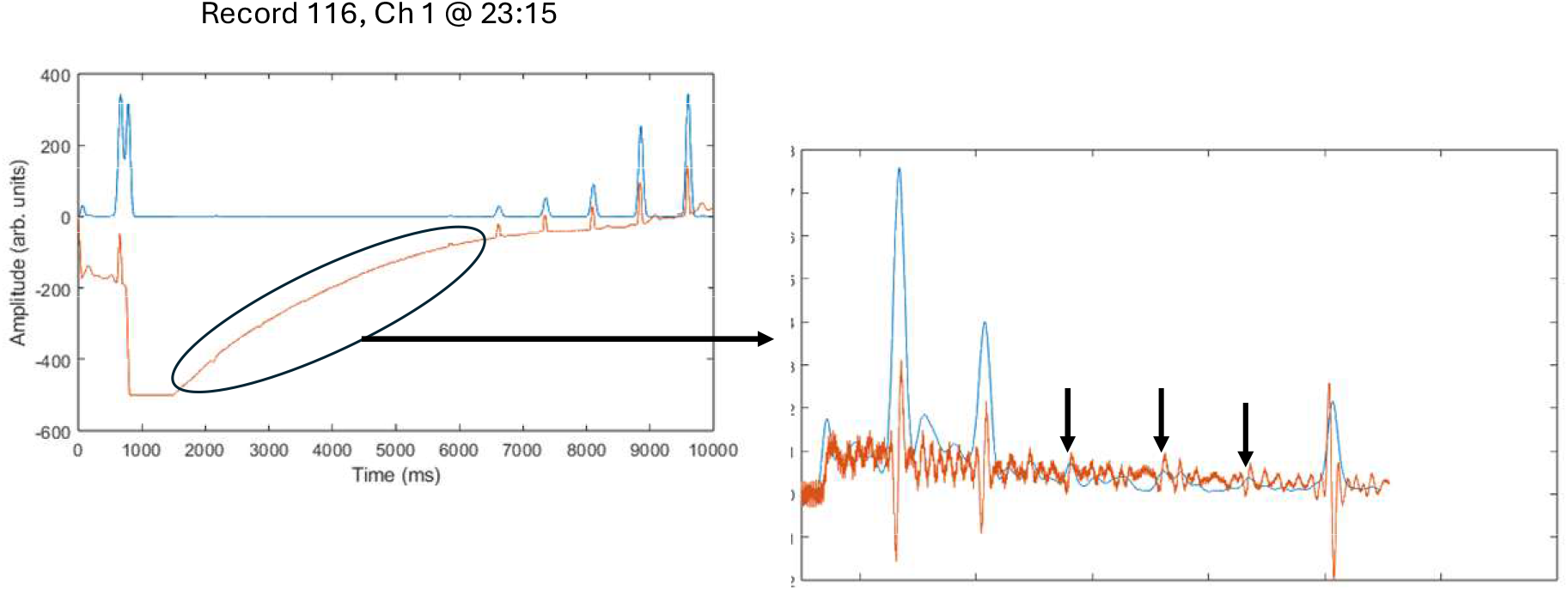

In the example shown in Figure 5, low quality true peaks in the dropout area (designated by arrows) are successfully detected because they form part of a highly structured sinus rhythm sequence.

### H. Supplementary Decision Rules

Approximately twenty rules, first order in 2-6 parameters, are applied after MLP scoring to reduce false positives and to moderately enhance sensitivity. Most of the rules involve 2-3 parameters selected from various combinations of MLP and other parameters (see table I). Examples of rules that reduce false positives by reducing (by at least 50%) scores of peaks to below detection threshold include:

- Peaks that are within a certain period (e.g. 350 ms) before or after a peak with a much larger Csig amplitude;
- Small Csig amplitude peaks that have a large ratio of the 5^th^ to 1^st^ largest deflections, indicative of sinusoidal noise;
- Peaks with a highly skewed ratio of largest up/down deflections.

Note that such sub-threshold peaks may still form part of structured sequences. (There is one segment level rule that modifies the peak shape up/down deflection rule: if a segment is low noise and peaks have high peak shape correlation, the rule is overridden; this allows detection of very odd peak shapes, e.g. the PVCs in the first few segments of Record 207.)

Rules that enhance sensitivity mainly involve locating relatively small amplitude peaks (in clean segments) that are followed by a T-wave, as determined by a slope histogram.

### I. Data Processing

The MLP training data was selected from approximately 70% of channel 1 signal segments. Given the highly skewed ratio of QRS to non-QRS peaks because channel 1 signals are generally very clean, the training set was selected so that it included at least 20% of non-QRS peaks. The supplementary decision rules were formed with respect to the entirety of channel 1 records.

Although care was taken to make these rules follow reasonable *a priori* assumptions, the algorithm must be considered overfit with respect to channel 1.

Ventricular flutter segments in Record 207 were excluded from the dataset.

All code was written in Matlab (The MathWorks, Inc., Natick, Massachusetts, United States), including the Deep Learning Toolbox.

## 3. Results

Sensitivity (SE) and Positive Predictive Value (PPV) for (overfit) channel 1 were 99.93% and 99.98% respectively. For channels 1 and 2 combined, SE and PPV were 99.96% and 99.98% respectively. Overall SE and PPV are less meaningful for channel 2 because it contains a number of noisy records. SE and PPV increased, as expected, with the exclusion of noisier records, as shown in the following table, which shows the percentage of channel 1 peaks that would qualify at a particular noise level.

### Sensitivity and PPV of Channel 2 results improve as noisier segments are excluded

**Sensitivity and PPV of Channel 2 results improve as noisier segments are excluded:**
| IBN Less Than | BWN Less Than | % Ch. 1 Peaks Qualify | CH 2 SE % | CH 2 PPV % |
| --- | --- | --- | --- | --- |
| 0.15 | 3.5 | 97.8 | 99.67 | 99.75 |
| 0.2 | 4 | 98.6 | 99.55 | 99.68 |
| 0.25 | 4.5 | 99.0 | 99.40 | 99.63 |

These results mask a great deal of inter-record variability because, even in channel 2, most of the records are very clean. Most of the losses in SE and PPV were due to the following records: 106, 200, 203, 208, 209, 221, and 231. <u>Excluding these six records and using the 0.15/3.5 noise criteria, SE and PPV were 99.90% and 99.92% respectively.</u> Note also that a threshold level was set for high frequency noise that excluded most of Record 101 for channel 2 but did not exclude any peaks from channel 1.

<u>Importantly, even in very noisy records, major rhythm features could still be tracked.</u> For example, in Record 106, channel 2 has many noisy signal segments interspersed with clean segments over approximately 6 consecutive minutes. (The MIT-BIH archive states that for channel 2, 13:41 out of 30:00 minutes are “noisy”; https://archive.physionet.org/physiobank/database/html/mitdbdir/records.htm#231, retrieved April 15, 2026.) Figure 6 shows an example of clean (channel 1) and noisy (channel 2) segments. Bigeminy is apparent from the channel 1 segment but is somewhat obscured by artifacts in the channel 2 segment. However, the structured sequence search feature of the present work enables the assessment of a bigeminal rhythm, as shown in the Poincare plot on the right side of the figure. The true RR interval pairs are indicated by red x’s while the detected RR pairs are indicated by blue circles. Despite the noisiness of the detected RR intervals, the bigeminal pattern, along with the presence of ventricular couples, is evident. Indeed, the structured search and scoring could be refined considering this large scale pattern and the noise of the detected RR intervals further reduced.

**Figure 6.**
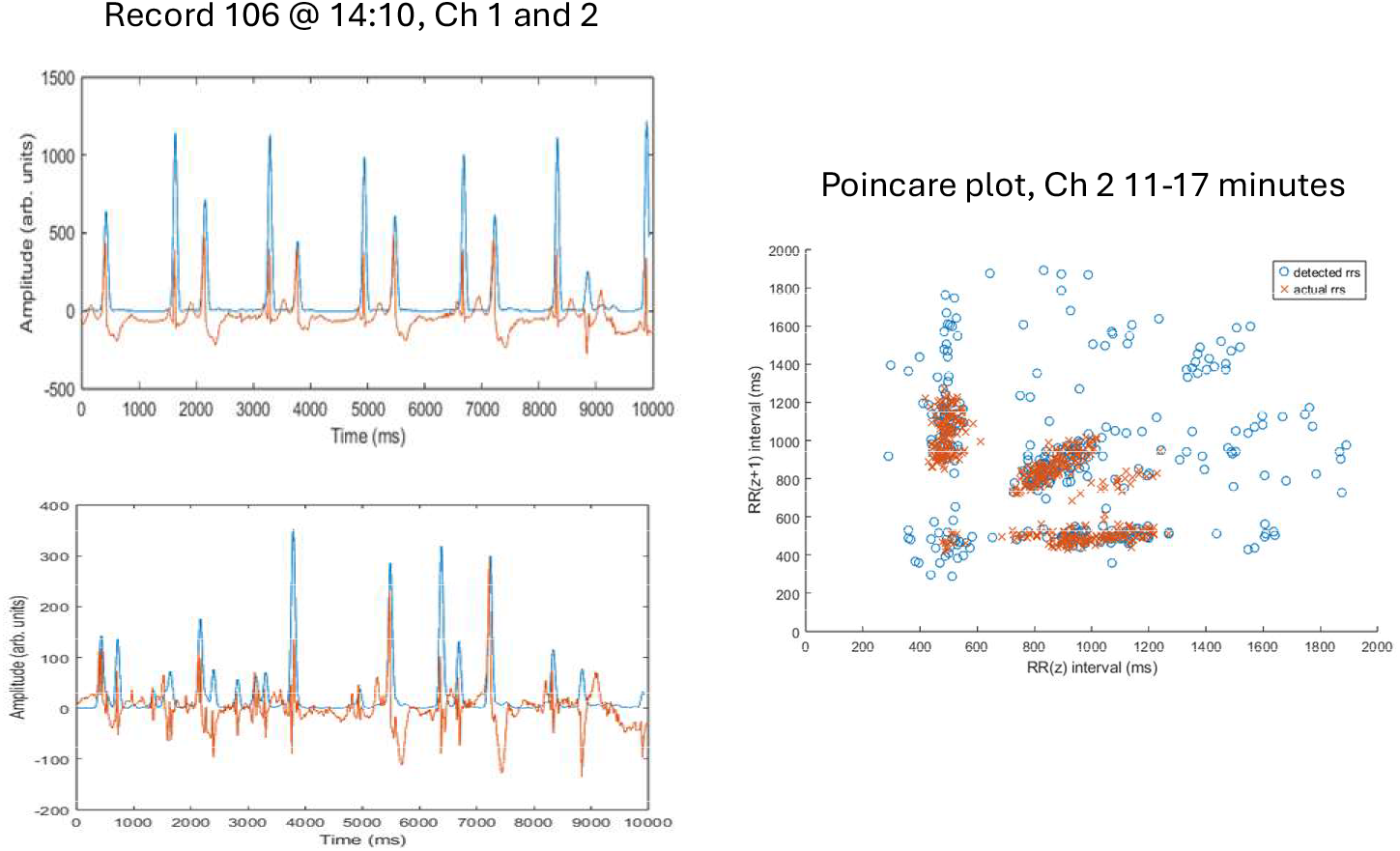

## 4. Discussion

The salient results of the present work are that combinatorial optimization enables a relatively small number of parameters to capture the detection information in the MIT-BIH DB, that appropriate noise measures correlate directly detection performance, and, relatedly, these same noise measures can produce very good results when used for multi-channel fusion. The present work also suggests that heart rhythm tracking may be possible even in relatively noisy signals that might be discarded by conventional ECG processing.

The multiple channel results in this work for the MIT-BIH (99.96% SE/99.98% PPV) compare to (99.94%/99.97%) for a superior single channel CNN[8], but since the supplementary decision rules were fashioned with regard to the entirety of the channel 1 dataset, this cannot be considered a fair contest. Regarding channel 2, the literature reveals few if any results for comparison.

The problems associated with the seven channel 2 records that degraded SE/PPV are summarized in the following table:

| Record | Issues |
| --- | --- |
| 106 | Noise measurement; small amplitude sinus QRS; structured search |
| 200 | Noise measurement |
| 203 | Relatively high amplitude F waves |
| 208 | Noise measurement; small amplitude sinus QRS; structured search |
| 209 | Noise measurement/insufficient selection of structured sequence |
| 221 | Small amplitude sinus QRS/over-filtering; T wave oversensing; QRS like artifacts |
| 231 | Heart block P-waves need to be removed (apart from record level designation which removes the problem in multi-channel fusion) |

Algorithmic improvements could alleviate many of these problems. Records 106, 208 and 221 exhibited similar signal patterns: small sinus QRS followed by large T waves and frequent large amplitude PVCs. These lead II patterns are quite different than the channel 1 V1 patterns which served as the basis for MLP training and rule choice. To perform properly in this situation, the algorithm’s T wave discrimination requires improvement. With regard to Record 231, P wave cancellation in the heart block context has been implemented but was developed with regard to this dataset so the benefit of this algorithmic feature is not reflected in the results of the present work.

Channel 2 of Record 208 contains a number of segments that would benefit from T wave tracking. Figure 7 shows filtered and unfiltered versions of two segments. As shown in the right hand side unfiltered version, the (sinus) QRS complexes are very small in relation to the following T waves; the few large peaks correspond to ventricular beats. Even after mild time domain smoothing, as shown in the bottom right panel, the QRS complexes are obliterated. Yet, such smoothing may be necessary if high frequency noise is present, as is the case in the left hand side segment. In this case, tracking T waves may enhance rhythm tracking and prevent noisy signal segments from either being discarded or result in an improper detection of bradycardia. This feature was not implemented but experimentation suggests that it is feasible.

**Figure 7.**
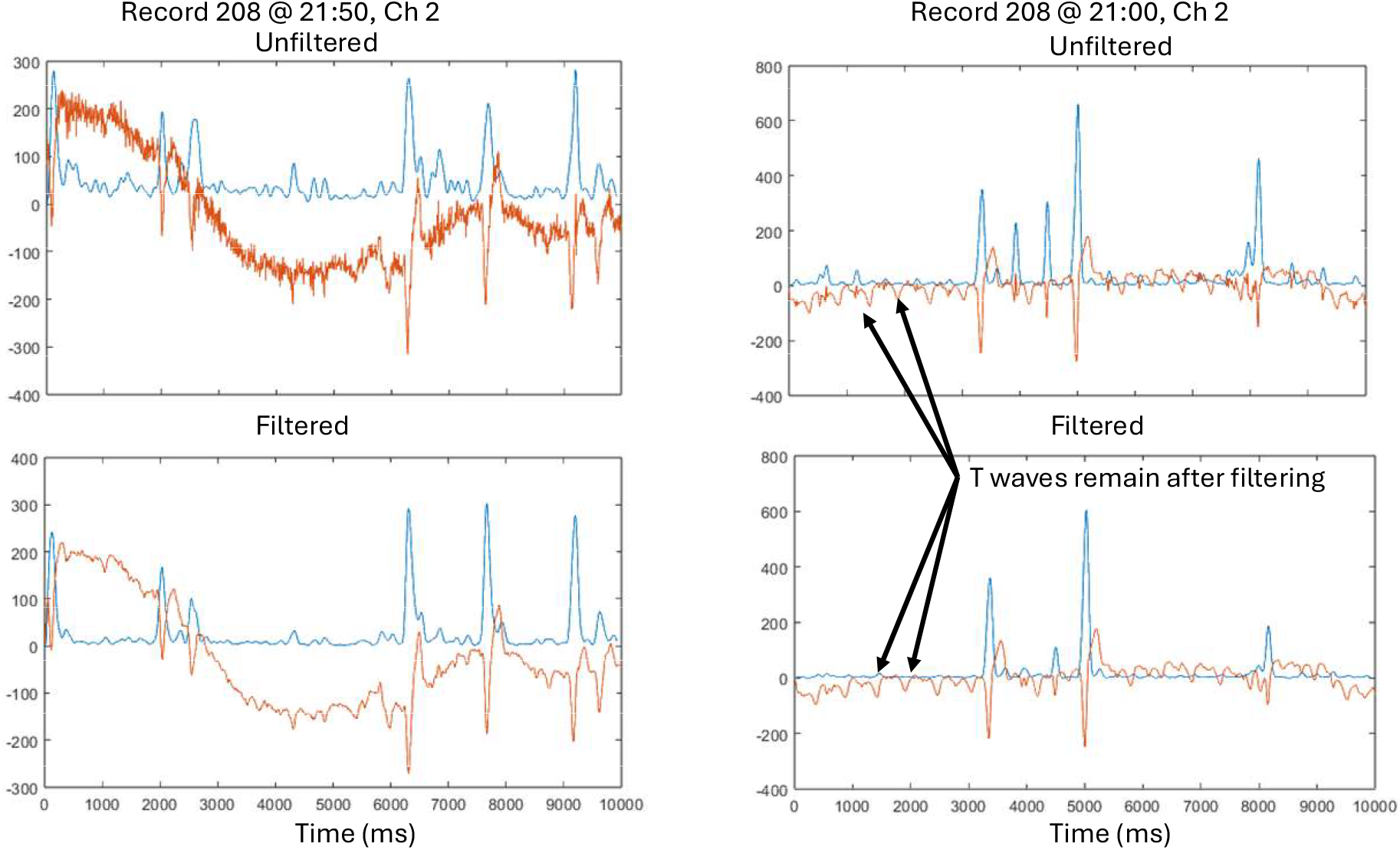

The heart block example described with reference to Figure 4 shows that, even beyond QRS detection, explicit analysis of temporal correlations can facilitate rhythm assessment. In particular, an analysis of the distribution of time between P waves and QRS complexes can help to distinguish between Mobitz types I and II heart block. Relatedly, the peak shape similarity measure, along with size and shape measures, can help to cluster different peak morphologies.

A combinatorial approach to QRS detection takes advantage of critical statistical information to enable superior noise resistance compared to “image recognition” type approaches. Specifically, this approach analyzes both the temporal and size/shape patterns of heartbeat components and <u>noise</u>. Although convolutional neural networks have been applied to QRS detection with impressive results[8,9], they are almost certainly inferior in the high noise environments that require explicit consideration of temporal correlations. Further, CNNs do not readily lend themselves to multiple channel fusion, tend to have poor temporal resolution, and cannot explicitly analyze important temporal correlations (e.g., PQ interval).

## 5. Conclusion

This work provides compelling evidence for the use of combinatorial optimization for QRS detection. This work also suggests that the discrete, parameter extraction and temporal correlation approach to peak analysis could greatly aid heart rhythm analysis and reduce false positive alerts [10] from automated ECG processing software. Because combinatorial optimization is such a broad methodology, it admits a large number of specific implementations that may involve machine learning techniques implemented in a hierarchical fashion. Much work remains to be done to achieve optimal analysis of the noisy ECG, which may become increasingly important in the age of wearables [11-13].

